# Disruption of the ecdysone oxidase gene *fezzik* drives transcriptomic, metabolic, and sex-specific lifespan changes in *Drosophila melanogaster*

**DOI:** 10.64898/2026.11.20.768145

**Authors:** Zhihui Bao, Amanda Glaser-Schmitt

## Abstract

Phenotypic variation within species is extensive. However, connecting it to underlying genetic variation remains an ongoing challenge, particularly for genes that influence complex traits subject to natural selection. Previous studies have detected selection on expression of the *Drosophila melanogaster* ecdysone oxidase gene *fezzik* (*fiz*), a systemic growth inhibitor that affects multiple phenotypes, although the molecular mechanisms through which *fiz* mediates its effects on phenotype remain largely unknown. Here, we performed RNA-seq, gene co-expression network analysis, and untargeted metabolomics in larvae with disrupted *fiz* expression to identify the systemic effects of *fiz* expression. 33 genes were consistently differentially expressed across two genetic backgrounds during *fiz* disruption, with *Chronologically inappropriate morphogenesis* (*chinmo*), a critical larval regulatory gene which was negatively associated with *fiz* expression, showing the largest and most consistent change. Overall, *fiz* expression was positively associated with expression of glycolysis-related genes and energy availability metabolites, but negatively associated with cell growth/proliferation and long chain fatty acids, which are important energy storage components. Thus, our results suggest that *fiz* is involved in energy metabolism, which is in line with its known effects on growth, body size, and malnutrition/starvation. These are likely mediated in part by dopamine signaling, as we detected decreased L-dopa levels during *fiz* disruption. L-dopa is also known to affect lifespan, likely playing a role in the sex-specific effect of *fiz* expression on lifespan we identified here. Interestingly, although their expression affected males similarly, *fiz* and *chinmo* expression affected female lifespan in opposing directions, suggesting that *chinmo* expression may partially mediate *fiz*’s effects on lifespan. By identifying the molecular mechanisms underlying a case of natural selection on a pleiotropic gene, this study sheds a deeper light on the evolution of complex traits and biodiversity, as well as the complex interaction between natural selection, genetic variation, and organismal phenotype.

**Author Summary:** A population’s history leaves distinctive patterns in the genome, allowing us to identify genes subject to different forms of selection. However, identifying the organismal trait under selection as well as the molecular mechanisms underlying changes during selection remains an ongoing challenge, especially when genes affect multiple complex traits. Understanding the mechanisms through which a selected gene affects phenotype can help us better understand the evolution of complex traits and how they contribute to phenotypic variation and the maintenance of biodiversity. Here, we use the fruit fly gene *fezzik* (*fiz*) to examine the molecular mechanisms underlying a case of complex selection on a gene affecting multiple complex traits. By examining gene expression and metabolites when *fiz* expression is disrupted, we found that *fiz* likely affects many traits by affecting the allocation of energy storage versus expenditure. These changes are most likely mediated in part by changes in dopamine signaling and expression of an important developmental regulatory gene. Based on our findings, we further identified lifespan as a novel trait that *fiz* influences in a sex-specific manner. Our study underscores the complexity of the interplay between selection, genetic variation, and organismal traits, including the underlying evolutionary and molecular mechanisms that shape them.

## Introduction

Connecting phenotype to genotype remains a complex, central challenge in genetics and evolutionary biology (reviewed in Martin et al. 2011, Gjuvsland et al. 2013, Tautz et al 2026). Understanding this connection is particularly important in cases of adaptation, where signs of selection can be detected at the DNA sequence level, but the phenotype under selection often remains difficult to determine. In non-model organisms or cases where the traits affected by a selected gene are unknown or myriad, making the connection between genotype and the phenotype under selection can be particularly challenging. However, elucidating the mechanisms of how selection shapes genetic variation and in turn phenotypic variation, especially in complex cases, is important for understanding the maintenance of biodiversity as well as the tempo and mode of evolution.

For example, human height shows extensive variation, is highly polygenic, and is thought to have been shaped by both adaptation and sexual selection (reviewed in Stulp and Barrett 2016). Studies have detected signs of polygenic selection for increased height in a UK population (Fields et al. 2016, Chen et al. 2020, but see Sohail et al. 2019), which has shed light on how selection occurs on complex traits. Another single gene example is the *Drosophila melanogaster* cytochrome P450 gene *CypCg1*, which is well-known for its resistance allele that underwent a selective sweep for resistance to the widespread use of the pesticide DDT in the 1940s and 1950s (Catania et al. 2004). However, in the absence of DDT, this gene has been subject to sexually antagonistic selection on the resistance and wild-type alleles driven by contrasting effects on fecundity, developmental time, and reproductive success between the sexes (Smith et al. 2011). Thus, the mechanisms that shape the interplay between selection and genetic and phenotypic variation can be complex and difficult to untangle.

In this study, we focus on *fezzik* (*fiz*), another *D. melanogaster* gene with a complicated selection history. Its protein sequence shows evidence for adaptive evolution (Langley et al., 2012; Saminadin-Peter et al. 2012), and expression of *fiz* is 2– 5-fold higher in cosmopolitan (non-sub-Saharan African) than in sub-Saharan (ancestral range) populations in both adult and larval stages (Meiklejohn et al. 2003, Hutter et al. 2008, Saminadin-Peter et al. 2012, Glaser-Schmitt et al. 2013, Glaser-Schmitt and Parsch 2018, 2023), which is driven by variation in an upstream *cis*-regulatory element (CRE). This CRE was the target of a selective sweep for increased *fiz* expression in cosmopolitan populations (Saminadin-Peter et al. 2012, Glaser-Schmitt et al. 2013) as the species expanded out of sub-Saharan Africa (Glaser-Schmitt and Parsch 2018). Although other SNPs are thought to be the original targets of the sweep (Glaser-Schmitt and Parsch 2018), one SNP drives the majority of this expression divergence (Glaser-Schmitt and Parsch 2018) and is thought to be maintained at intermediate frequency in cosmopolitan populations by a combination of sexually antagonistic and temporally fluctuating selection (Glaser-Schmitt et al. 2021, 2024a).

When signs of positive selection were first identified in its CRE, *fiz*’s function and therefore the phenotype under selection were unknown (Saminadin-Peter et al. 2012, Glaser-Schmitt et al. 2013). Since then, *fiz* expression has been found to affect body size and proportion, larval growth rate, starvation resistance, as well as cold and insecticide tolerance (Glaser-Schmitt and Parsch 2018, Glaser–Schmitt et al 2021). It acts as a systemic growth inhibitor (Glaser-Schmitt and Parsch 2018), and decreased *fiz* expression during larval malnutrition or imbalanced diet results in faster larval growth, shorter development time, and larger body size (Grenier et al. 2023, Cavigliasso et al. 2024). The fiz protein has ecdysone oxidase activity, acting on both ecdysone and 20-Hydroxyecdysone (Cavigliasso et al. 2024), which is in line with the idea that *fiz* affects growth and body size via a modulation of ecdysone signaling (Glaser-Schmitt and Parsch 2018). However, the precise molecular mechanisms through which *fiz* expression affects these final organismal traits remain unknown, which what this study aims to identify.

High-throughput RNA sequencing (RNA-seq) in individual tissues can help identify how expression contributes to complex adaptive phenotypes (reviewed in Hoedjes et al. 2025), such as those segregating among *D. melanogaster* populations (Durmaz Mitchell et al 2025). Therefore, we used RNA-seq in two genetic backgrounds to identify how systemic disruption of *fiz* expression affects transcription and gene co-expression networks in the larval fat body, which is the *D. melanogaster* equivalent of mammalian liver and white adipose tissue and is critical for a myriad of functions potentially related to phenotypes that *fiz* expression affects, such as developmental timing, growth coordination, energy storage and metabolism, the immune response, detoxification, and nutrient sensing (Mirth and Riddiford 2007, Liu et al 2009, Arrese and Soulages 2010, Ǫuinn et al. 2012, Mirth and Shingleton 2012). Moreover, *fiz* expression is enriched (Krause et al 2022) and shows divergent expression between a derived and ancestral population (Glaser-Schmitt and Parsch 2023) in this tissue. To better connect the identified expression changes to final organismal phenotype, we further performed untargeted metabolomics during *fiz* expression disruption. Our expression and metabolomics analyses suggest that *fiz* is involved in energy metabolism with higher *fiz* expression associated with more energy free for expenditure, but lower *fiz* expression associated with increased growth and energy storage. Moreover, these effects are likely modulated in part by changes in dopamine signaling and expression of *chronologically inappropriate morphogenesis* (*chinmo*), an important developmental regulatory gene, suggesting *fiz* affects phenotype via multiple signaling cascades. Our approach further suggested a new phenotype that *fiz* expression affects, lifespan, which we confirmed is affected by *fiz* expression in a sex-specific manner, suggesting that it may be connected to trade-offs related to previously identified sexually antagonistic and temporally varying selection acting on *fiz*’s upstream CRE (Glaser-Schmitt et al 2021, 2024a).

## Results

To understand how disruption of *fiz* expression affects the expression of the rest of the transcriptome, we performed RNA-seq in the larval fat body in a control strain (controlKD) and one in which *fiz* expression was knocked down (*fiz*KD) via RNAi (see Materials and Methods for more details). Knockdown of *fiz* in the in *fiz*KD strain was approximately 2-fold (-1.086 log_2_-foldchange–L2FC; Table S1) and *fiz*KD samples clustered separately from controlKD samples in a principal component analysis (PCA; Fig 1A). While RNAi in an otherwise homogeneous genetic background is an excellent tool to interrogate the contribution of a gene’s expression to phenotype (reviewed in Heigwer et al 2018), the more heterogenous genetic backgrounds that are present in natural populations (Kapun et al 2021, Nunez et al 2025) can lead to genetic background effects on expression (Osada et al 2017, Glaser-Schmitt et al 2018, GTEx Consortium 2020, Puixeu et al 2023, Glaser-Schmitt et al 2024b) and phenotype (Garcia et al 2020, Glaser-Schmitt et al 2021, Vea et al 2026). To better understand how disruption of *fiz* expression affects the rest of the transcriptome across multiple genetic contexts, including natural populations, we further analyzed our knockdown data in conjunction with RNA-seq data in 5 strains from a natural European population (Glaser-Schmitt and Parsch 2023), one of which contained a *fiz* hypomorph allele (Glaser-Schmitt and Parsch 2018). In this combined dataset including natural population and lab backgrounds, disruption of *fiz* was approximately 6-fold (-2.57 L2FC; Table S1) across strains in which *fiz* expression had been disrupted (*fiz*DIS), and after batch correction the *fiz*DIS strains clustered together (Materials and Methods; see Fig S1 for PCA before batch correction), although the control strain (controlDIS) cluster slightly overlapped (Fig 1B).

**Fig 1:**
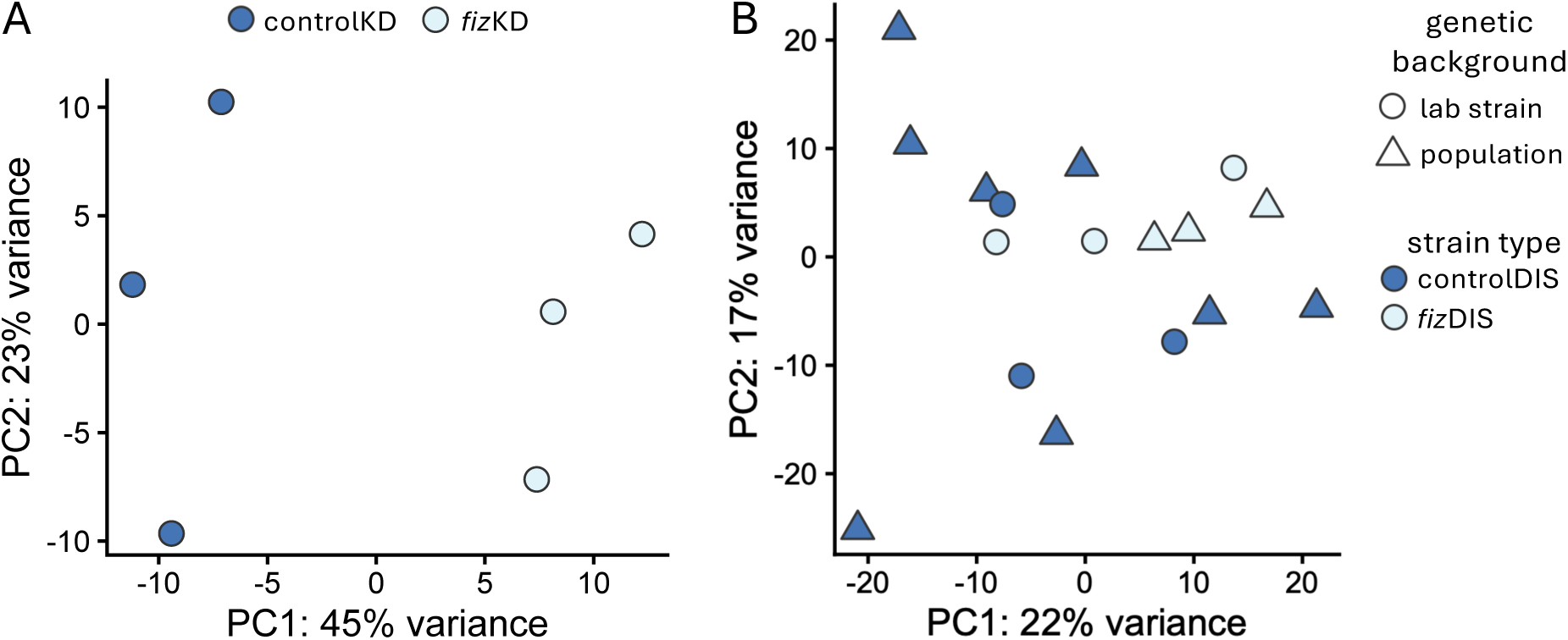
PCA of gene expression profiles in A) *fiz*KD (light blue) and controlKD (dark blue) and B) *fiz*DIS (light blue) and controlDIS (dark blue) strains. Shape indicates the genetic background.

### Metabolism-, especially glycolysis-related genes, were consistently down-regulated during *fiz* expression disruption

We used DESeq2 (Love et al. 2014) to identify protein coding and non-coding genes (ncRNAs) differentially expressed (DE) when *fiz* expression was disrupted. When we knocked down *fiz* expression in the lab strain background, 1.21% and 1.26% of genes were up- and down-regulated in the *fiz*KD strain, respectively (212 genes in total; Table 1). In keeping with the known effects of *fiz* expression on developmental time and body size (Glaser-Schmitt and Parsch 2018), genes up-regulated in the *fiz*KD strain were enriched for biological process GO terms related to regulation of anatomical structure size as well as development, morphogenesis, and signaling processes but no KEGG pathways (Fig S2 and Table S2). Down-regulated genes were enriched for the small molecule catabolic process GO term and the galactose as well as sucrose and starch metabolism pathways (Fig S2 and Table S2). Similar to results in just the lab background, when we examined the full dataset, we detected 1.45% and 0.89% of genes as up- and down-regulated in the *fiz*DIS strains, respectively (211 genes in total; Table 1). Down-regulated genes in *fiz*DIS strains were enriched for GO terms related to glycolytic and catabolic processes as well as pathways related to glycolysis and glucogenesis (Fig S2 and Table S3), which suggests that normal *fiz* expression is associated with the conversion of glucose into energy rather than energy storage, which is consistent with *fiz*’s effects on growth and body size (Glaser-Schmitt and Parsch 2018; Grenier et al. 2023; Cavigliasso et al. 2024). Up-regulated genes were more dissimilar to results using just the lab background and were enriched for PIWI-interacting RNA (piRNA) and regulatory ncRNA processing as well as mitosis and chromosome condensation and segregation related GO terms and pathways involved in DNA replication and repair (Fig S2 and Table S3). Thus, genes involved in glycolysis and metabolism more generally were positively associated with *fiz* expression (i.e. were down-regulated when *fiz* expression was disrupted), but genes negatively associated with *fiz* expression (i.e. were up-regulated when *fiz* expression was disrupted) were more diverse among genetic backgrounds.

**Table 1:** DE genes when *fiz* expression is disrupted**Fig 2**: Genes DE in the A) *fiz*KD and B) *fiz*DIS genetic backgrounds. Significantly DE genes (see Methods) are shown in blue. The gene *chinmo* was the most highly DE in both backgrounds.

|  | *fiz*KD | *fiz*DIS | Shared | only KD | only DIS |
|---|---|---|---|---|---|
| Genes | 8,566 | 9,015 | 8,510 | 56 | 505 |
| Up (%) | 104 (1.21) | 131 (1.45) | 20 (0.24) | 84 | 111 |
| Down (%) | 108 (1.26) | 80 (0.89) | 13 (0.15) | 95 | 67 |

Shown are the number of expressed genes as well as the number and percentage (in parentheses) of up- or down-regulated genes in strains in which *fiz* expression is disrupted.

### Only 33 genes were consistently DE when *fiz* expression is disrupted

To identify which genes were consistently DE when *fiz* expression was disrupted, we examined the overlap between the RNAi and the full dataset. The overlap in disrupted genes between the lab background alone and the analysis with two genetic backgrounds was very low, with only 33 genes shared in total, representing 9.30% and 7.43% of all up- and down-regulated genes being shared, respectively (Table 1). This low overlap suggests that outside of a few key genes, the genes affected when *fiz* expression is disrupted is largely genetic background-specific, despite all *fiz*DIS flies displaying similar final organismal phenotypes in comparison to control strains (Glaser-Schmitt and Parsch 2018). Thus, it may be that *fiz* expression affects downstream organismal phenotype via consistent changes in the expression of only a small number of genes. In keeping with this interpretation, only 4 genes showed a consistent L2FC of 2 or greater (Table S4) in the lab background: *hulk* (*hlk*), *nirvana 3* (*nrv3*), *chinmo*, which were up-regulated in the *fiz*KD line, and *Tetraspanin 2CA* (*Tsp2CA*), which was down-regulated (Table S4); while only one gene, *chinmo*, met this threshold and was up-regulated in *fiz*DIS flies (Fig 2, Table S4). This *chinmo* up-regulation occurred in both sexes, but was weaker in males (Fig S3). Although genes meeting this DE threshold represent our best candidates for genes that modulate *fiz*’s effects on phenotype, other shared genes are also likely to contribute. For example, *Frost* (Table S4) is known to affect cold tolerance (Colinet et al 2010, Garcia et al 2020), which likely plays a role in *fiz*’s sex-specific effect on cold tolerance (Glaser-Schmitt and Parsch 2018).

**Fig 2:**
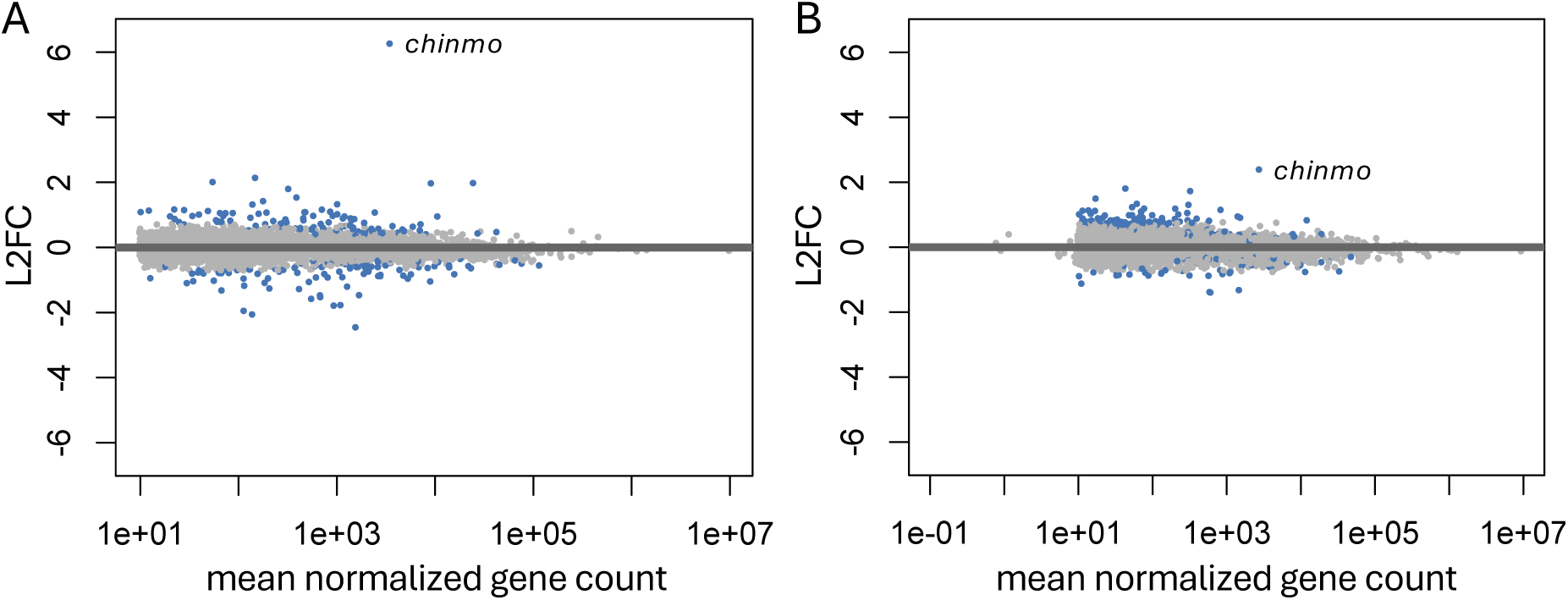
Genes DE in the A) *fiz*KD and B) *fiz*DIS genetic backgrounds. Significantly DE genes (see Methods) are shown in blue. The gene *chinmo* was the most highly DE in both backgrounds.

Of the shared 33 genes, 6 (18.18%) were non-coding (Table S4), suggesting that *fiz* expression also affects non-coding gene expression, which is in line with our finding of enrichment for genes with GO terms related to regulatory ncRNA processing among up-regulated genes in the full dataset (Fig S2 and Table S3). When we ran a GO enrichment analysis for these shared genes, we did not detect an enrichment of any GO terms or pathways among up-regulated genes, which is unsurprising given their small number. Similar to analyses using the full and just the RNAi dataset, we detected an enrichment of genes with GO terms related to glycolysis and catabolism as well as the glycolysis/gluconeogenesis pathway (Fig S4 and Table S5) among down-regulated genes. Thus, less than 0.4% of genes were consistently DE across genetic backgrounds when *fiz* expression was disrupted, and the down-regulated genes showed a similar enrichment for metabolism and glycolysis-related processes, suggesting that *fiz* may play a role in energy metabolism. In line with this interpretation, we detected *Coenzyme ǪC* (*CoqC*) among the shared up-regulated genes (Table S4). *CoqC* is an important component in Coenzyme Ǫ biosynthesis (Öztürk-Çolak et al 2024), which is crucial for the mitochondrial electron transport chain and energy metabolism (reviewed in Wang et al 2024b).

### *fiz* expression-associated co-expression modules lacked DE genes

In order to better understand how *fiz* expression affects gene co-expression networks, we used our full dataset to perform a weighted gene co-expression network analysis (WGCNA; Langfelder and Horvath 2008) in which we clustered genes into modules with correlated gene expression and identified 6 co-expression modules. The majority of analyzed genes (94.9%) were not assigned to a specific co-expression module (grey module; Fig 3A), while the rest were assigned to five modules 33–156 genes in size (Fig 3A). Interestingly, all genes we identified as having consistent expression differences when *fiz* expression is disrupted belonged to the unassigned, grey module, which underscores that outside a few, key DE genes, the downstream effects of *fiz* expression on the transcriptome may be genetic background-specific.

**Fig 3:**
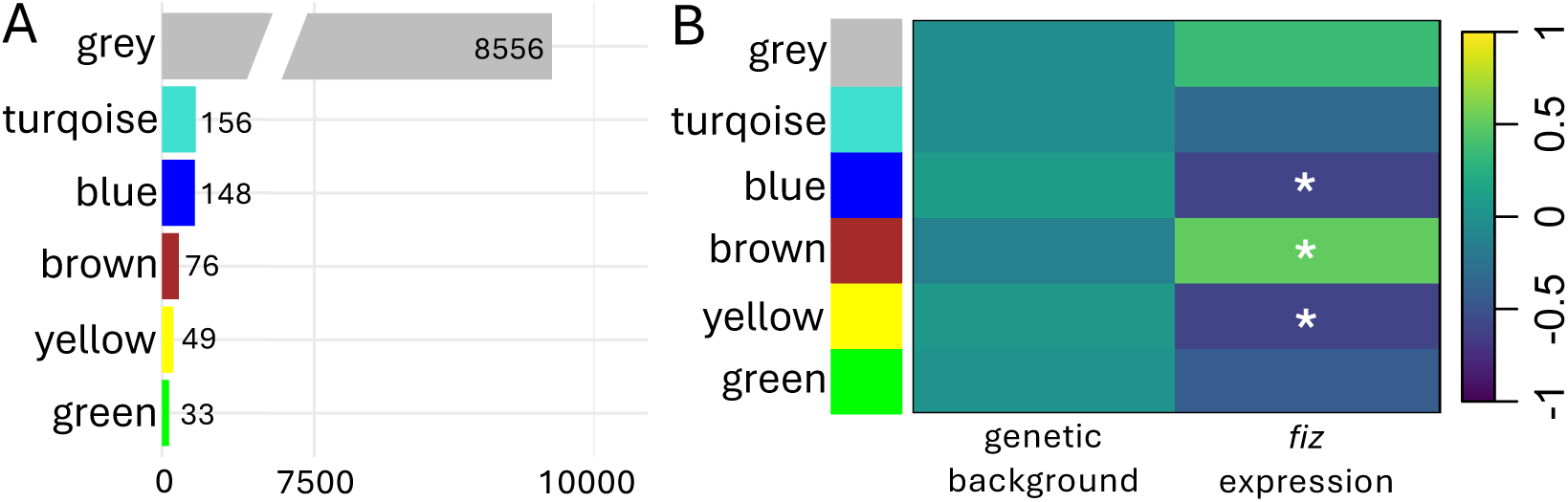
WGCNA revealed A) five co-expression modules, B) three of which were significantly associated with *fiz* expression. Colors correspond to module names. A) Number of genes belonging to each module is shown. B) Color indicates the strength of association between a module and genetic background or *fiz* expression. The grey module consists of genes that could not be classified into the other modules. Significance was assessed using the student asymptotic *P*-value from the Pearson’s correlation. \**P* < 0.05

However, when we correlated the module eigengenes (i.e. the module summaries) with genetic background (see Methods), we did not detect any significant associations between any modules and genetic background (Fig 3B), suggesting that any genetic background effects occur on an individual gene basis rather than through correlated expression changes in consistent suites of genes.

Next, we correlated the module eigengenes with *fiz* expression and identified 3 modules (brown, blue, and yellow) as associated with *fiz* expression (Fig 3B), for which we performed GO biological process and KEGG pathway enrichment analyses. The brown module, which was positively associated with *fiz* expression (Fig 3), was enriched for protein catabolic biological processes and proteasome pathways (Fig S5 and Table S6). The blue module, which was negatively associated with *fiz* expression (Fig 3), was enriched for biological processes related to mitosis and chromosome segregation and organization as well as DNA replication pathways (Fig S5 and Table S6), which are all important for the cell division required during growth, in keeping with *fiz*’s known action in growth suppression (Glaser-Schmitt and Parsch 2018; Grenier et al. 2023; Cavigliasso et al. 2024). The yellow module, which was also negatively associated with *fiz* expression (Fig 3), had no enriched GO terms but was enriched for xenobiotic metabolism pathways (Fig S5 and Table S6), which may be related to increased pesticide resistance when *fiz* expression is disrupted (Glaser-Schmitt and Parsch 2018). Retinol metabolism and carbohydrate metabolism, specifically ascorbate and aldarate metabolism, pathways were also enriched (Fig S5 and Table S6), alterations in the expression of both of which have been linked to changes in lifespan (Moskalev et al 2019; Wang et al 2024a). Thus, our WGNCA detected modules of co-expressed genes both significantly associated with *fiz* expression and connected to known and potential new phenotypes that *fiz* expression affects, despite these modules containing no DE genes.

### *fiz* expression affects dopamine signaling and energy storage/availability

Results from our WGNCA and RNA-seq analyses suggested that *fiz* expression may affect metabolism (Figs S2 and S5, Tables S2–3 and S5–6). We therefore performed untargeted metabolomics using liquid chromatography-mass spectrometry (LC-MS) to identify DE metabolites between whole *fiz*KD and controlKD larvae (see Methods for more details). Of the 2,881 positive ion-mode and 858 negative ion-mode metabolites detected, 36 (30 and 6, respectively) were down-regulated in the *fiz*KD strain, while 45 (14 and 31, respectively) were up-regulated (Table S7). To better understand the pathways these metabolites are involved in, we performed a KEGG pathway enrichment for DE metabolites detected in positive and negative ion-mode separately. Negative ion-mode DE metabolites were enriched for terpenoid backbone biosynthesis (Table S8), which occurs in plants and is unlikely to be related to *fiz* expression. On the other hand, positive ion-mode DE metabolites were enriched for tyrosine metabolism (Table S8), which plays an import role in metabolic homeostasis (Kosakamoto et al 2024) and is associated with variation in lifespan (Parkhitko et al 2020). Although we did not detect any enrichment for carbohydrate metabolism or glycolysis as we did in our WGCNA and RNA-seq analyses (Figs S2 and S5, Tables S2–3 and S5–6), we did detect a 6-glucose derivative as down-regulated in the *fiz*KD strain, albeit at the lowest confidence level (level 3, see Methods; Table S7).

We next focused on the 18 of 81 DE metabolites (9 up- and 9 down-regulated; 22.2%) that were identified at the highest confidence level (level 1; see Methods). 4 out of 5 DE lipids, including all 3 DE long chain fatty acids, which are essential for energy storage in the form of triglycerides and also function as building blocks for membrane lipids and signaling molecules (reviewed in Heier and Kühnlein 2018), were up-regulated in the *fiz*KD strain (Table S7, Figs 4, S6). 1-tridecanoyl-sn-glycero-3-phosphoethanolamine (1-tri-sn-glyc-3-phosph), which is a glycerophosphoethanolamine, structural components of the plasma membrane that influence shape (van Meer et al 2008), was also up-regulated (Fig 4, Table S7). Also up- regulated was myristoylglycine (Fig 4, Table S7), a myristolyzed form of the amino acid glycine that plays crucial roles in cellular signaling, protein interactions, and membrane targeting when incorporated into the N-terminal end of a protein (Farazi et al 2001).

**Fig 4:**
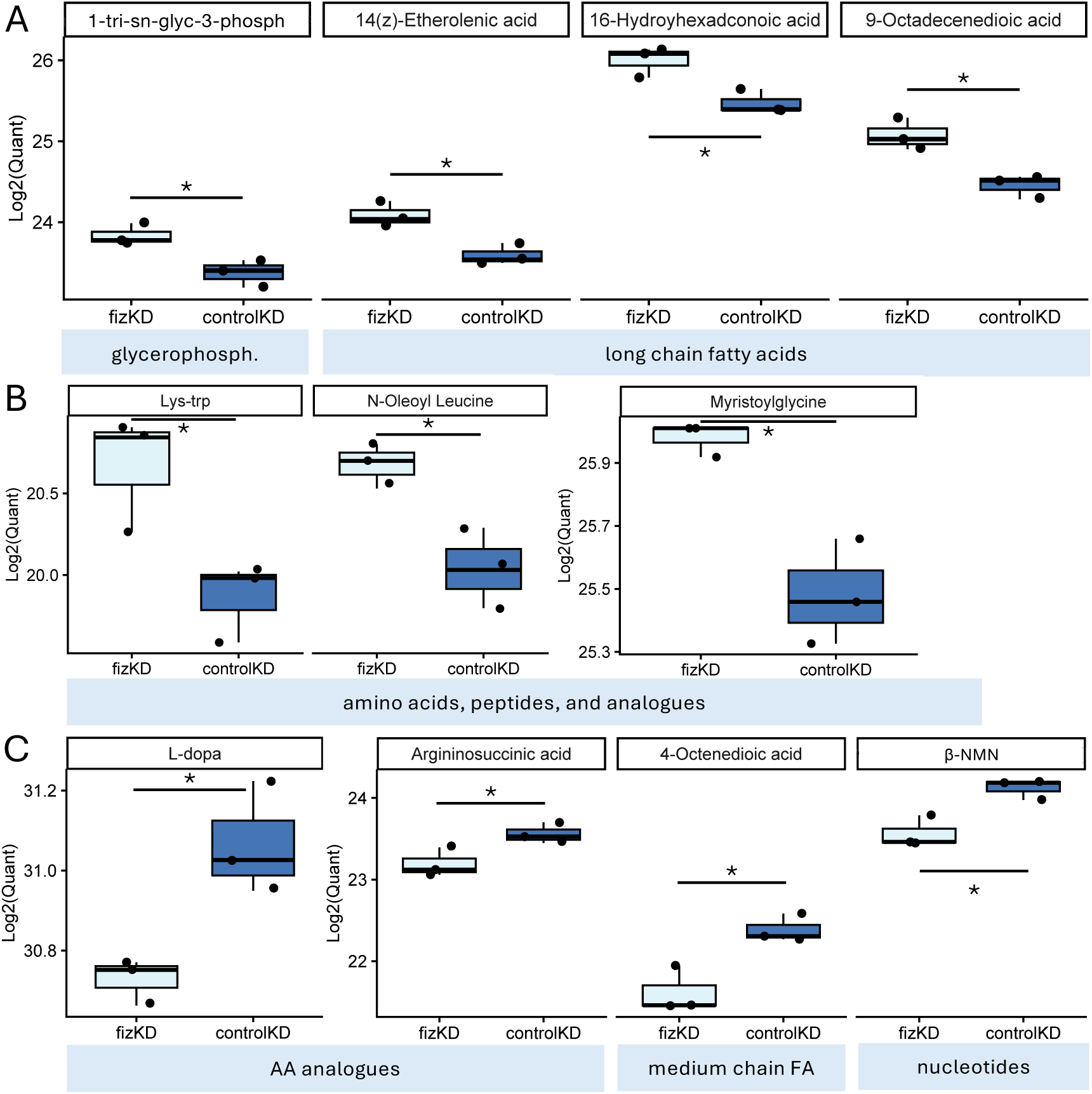
Levels, expressed in quantitative values (quant), of high confidence metabolites A,B) up- or C) down-regulated in *fiz*KD (light) in comparison to controlKD (dark) strains. Shown are DE A,C) lipids, including long or medium chain fatty acids (FAs) and glycerophospholipids (glycerophosph.), B,C) amino acids (AAs) or peptides and derivatives, and C) nucleotides. Significance was assessed with a students *t*-test. * *P* < 0.05.

Amino acids can serve as protein synthesis building blocks or cues during nutrient sensing in the fat body, serving as key regulators of systemic growth within the fat body (Manière et al 2020, Gillette et al 2021). Several other amino acids were also up- regulated (Fig 4, Table S7): N-oleoyl leucine, a derivative of the amino acid leucine, and Lys-Tryp, a dipeptide consisting of the amino acids lysine and tryptophan. Thus, metabolites negatively associated with *fiz* expression functioned in lipid energy storage and cellular functions crucial during growth (i.e. signaling and structural membrane components as well as protein synthesis).

Among down-regulated metabolites, we detected 4-octenedioic acid (Table S7, Fig 4), the only DE medium chain fatty acid, a type of fatty acid that can be quickly converted into energy and plays a role in regulating glucose and lipid metabolism (reviewed in Huang et al 2021). Also down-regulated was the arginine precursor, argininosuccinic acid, also known as argininosuccinate (Table S7, Fig 4), which is part of the urea cycle. Knockdown of genes involved in argininosuccinate metabolism results in increased triglyceride levels as well as decreased cold tolerance and lifespan (Heinrichsen et al 2014). L-dopa, also known as levodopa, a dopamine precursor derived from L-tyrosine, was also down-regulated (Table S7, Fig 4). In *Drosophila*, among other phenotypes such as sleep/arousal, sensory and larval exploratory behaviors (Neckameyer 1996, Andretic et al 2005), dopamine signaling is known to modulate the metabolic rate (Ueno et al 2012); while L-dopa, specifically, has been linked to metabolism-driven changes in lifespan (Wang et al 2024a) as well as changes in feeding behavior, body weight and lipid content (Bobrovskikh et al 2026).

Interestingly, two other L-dopa derivatives detected at a lower confidence level (level 3; see Methods) were also down-regulated (Table S7), further underscoring the down- regulation of dopamine signaling in the *fiz*KD strain. β-nicotinamide mononucleotide (β- NMN), the biologically active form of NMN, a precursor of the redox coenzyme nicotinamide adenine dinucleotide (NAD), which is critical for cellular redox homeostasis and metabolism (Xiao et al 2018) and is associated with variation in lifespan (Yadav et al 2026), was also down-regulated (Table S7, Fig 4). Increased NMN levels have been shown to increase levels of NAD+ (the oxidized form) specifically as well as decrease the NADH/NAD+ ratio (Yadav et al 2026, reviewed in Xie et al 2020), while increased NADH/NAD+ ratios and reduced NAD+ levels have been linked to reduced energy expenditure and obesity (reviewed in Xie et al 2020). Thus, metabolites positively associated with *fiz* expression were involved in dopamine signaling, lifespan, reduced lipid storage, and energy expenditure.

### *fiz* expression affects lifespan in a sex-specific manner

Results from our WGNCA and metabolomics analyses suggested that *fiz* expression might affect lifespan. Moreover, *chinmo*, which was up-regulated during disruption of *fiz* expression (Figs 2 and S3, Table S4), has been found to contribute to extended lifespan during dietary restriction (Pandey et al 2021). We therefore measured lifespan in *fiz*KD and controlKD flies as well as flies in which *chinmo* expression had been knocked down (*chinmo*KD) and the respective control line (controlSH, see Methods). When we knocked down *chinmo* expression, lifespan significantly decreased in both sexes (Wald test, *P* < 0.02 for both, Fig 5). Similarly, in males, knocking down *fiz* expression significantly decreased lifespan *(P* = 0.0321, Fig 5). However, we detected the opposite effect in females: knocking down *fiz* expression significantly increased lifespan (*P* = 0.004, Fig 5), an oppositional effect we would expect if *chinmo* and *fiz* expression are negatively associated and both genes affect lifespan. Thus, *fiz* expression affects longevity in a sex-specific matter, with female expression affecting lifespan in the opposite direction of *chinmo* expression.

**Fig 5:**
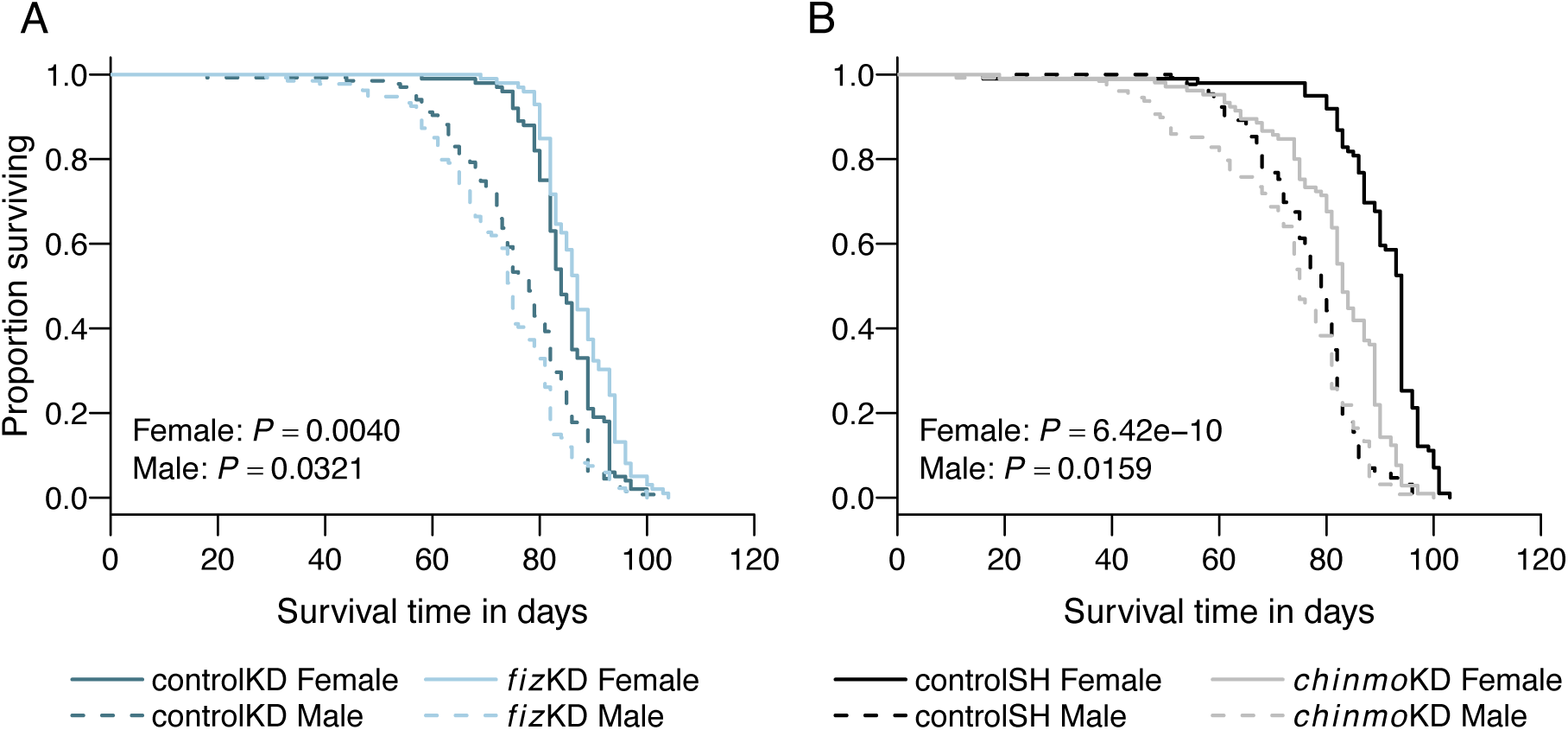
The effect of *fiz and chinmo* expression on lifespan. Kaplan–Meier curves show survival of A) *fiz*KD flies and controlKD and B) *chinmo*KD flies and controlSH at 25°C. Controls and RNAi flies are shown in A) dark and light blue and B) black and gray, respectively. Solid and dashed lines indicate females and males, respectively.

Significance was assessed with a Wald test comparing each RNAi genotype with its corresponding control using a Cox proportional hazards model separately for each sex and adjusting for collection batch (see Methods).

## Discussion

Taken together, our results suggest that *fiz* expression plays a role in energy metabolism: At normal or higher *fiz* expression levels energy is more readily available for immediate expenditure via the NAD+ precursor β-NMN or medium chain fatty acids and the expression of glycolysis-related genes (Fig 4, Tables S2–3, S5), whereas when *fiz* expression is lower or disrupted, energy is shuttled toward more long term energy storage via long chain fatty acids and basic cellular functions promoting cell growth and proliferation (Fig 4, Tables S2–3, S5). Maintaining the balance between energy expenditure versus storage is complex, requiring the integration of external and nutritional cues, signaling pathways, and individual tissues with each other and the organism as a whole, and this process is particularly important during growth and development (reviewed in Gillette et al 2021). Moreover, the larval fat body, where we identified differential transcription in this study, plays an important part in these processes as it is the main regulator of systemic growth and metabolism (Tennessen and Thummel 2011; Huang and Perrimon 2026) and is central to sensing nutritional signals including in response to metabolic status (reviewed in Tennessen and Thummel 2011; Huang and Perrimon 2026).

Outside of the DE genes and metabolites that we identified in this study (Figs 2, 4 and Tables S4, S7), the precise mechanisms through which *fiz* interacts with other components of energy metabolism remain relatively unknown. However, *fiz* is annotated as a glucose-methanol-choline oxidoreductase (Öztürk-Çolak et al 2024) and displays ecdysone oxidase activity acting on both ecdysone and 20- hydroxyecdysone (Cavigliasso et al. 2024). Ecdysone signaling is known to regulate triacylglycerol metabolism and lipid accumulation (reviewed in Heier and Kühnlein 2018). It has also been connected it to the regulation of energy metabolism and homeostasis (Homem et al 2014, Nishimura 2020). Although we did not detect any ecdysone derivatives among our DE metabolites (Table S7), ecdysone is known to play different roles and occur at different levels across life stages and tissues (reviewed in Boulan and Mirth 2026), which may have masked differences in ecdysone levels as we used whole larvae in our metabolomics analyses. In keeping with this interpretation, levels of active ecdysone, as approximated by *E74B* expression, have been shown to decrease when *fiz* expression in disrupted (Glaser-Schmitt and Parsch 2018). Thus, *fiz* likely modulates energy metabolism at least in part via its ecdysone oxidase activity and ecdysone signaling.

A role in energy metabolism is in line with *fiz*’s known action as a systemic growth inhibitor (Glaser-Schmitt and Parsch 2018; Grenier et al. 2023; Cavigliasso et al. 2024), with flies in which *fiz* expression has decreased or been disrupted displaying increased growth and larger body size. This effect on systemic growth is most likely mediated in part by *chinmo*, a BTB-zinc finger transcription factor and the most consistent and highly DE gene we detected, which was up-regulated during *fiz* disruption (Figs 2, S3, Table S4) and is known to promote larval growth (Chafino et al 2023). It is also likely mediated in part by *CoqC*, which is involved in Coenzyme Ǫ biosynthesis and therefore the electron transport chain (reviewed in Wang et al 2024b). *CoqC* expression was negatively associated with *fiz* expression (Table S4) and reduced electron transport chain activity severely curtails growth (Meiklejohn et al 2013). The increased body size is likely due at least in part to an accumulation of energy storage in the form of lipids, specifically long chain fatty acids, which increase as *fiz* expression decreases (Fig 4, Table S7). Moreover, this increased stored lipid content is likely due at least in part to *fiz*’s action on ecdysone (Cavigliasso et al. 2024), which regulates lipid accumulation (reviewed in Heier and Kühnlein 2018), as well as changes in *chinmo* expression in the fat body, where it is known to modulate fat metabolism (Pandey et al 2021). It is also most likely mediated at least in part by dopamine signaling as levels of L-dopa, which is commonly used experimentally to increase dopamine levels and signaling (for example in Bobrovskikh et al 2026), were positively associated with *fiz* expression. Dopamine levels have been shown to be negatively associated with total lipid content and body weight, which, interestingly, co-occurred with increased food intake (Bobrovskikh et al 2026), suggesting the *fiz*’s effects on growth and energy storage might also be mediated by nutritional intake. Indeed, dopamine signaling is known to inhibit foraging behavior and promote initiation of feeding (reviewed in Pool and Scott 2014). In keeping with this possibility, we detected several amino acids and a dipeptide, which can act as sensory cues and key regulators of systemic growth in response to nutritional input within the fat body (Manière et al 2020, Gillette et al 2021), as up-regulated when *fiz* expression was disrupted (Fig 4).

The increased energy reserves when *fiz* expression is decreased or disrupted likely contribute to *fiz*’s known effects on larval malnutrition (Kawecki et al 2021, Cavigliasso et al. 2024) and adult starvation resistance (Glaser-Schmitt et al 2021), the second of which is sex-specific (Glaser-Schmitt et al 2021). Interestingly, we found that the effect of *fiz* expression on lifespan was also sex-specific (Fig 5). Traits that affect fitness, and life history traits such as lifespan in particular, are often correlated, resulting in trade-offs between different components of fitness (reviewed in Flatt 2020). Starvation resistance is thought to be the target of sexually antagonistic and temporally fluctuating selection acting on *fiz*’s upstream CRE (Glaser-Schmitt et al. 2021).

However, the sex-specific effect of *fiz* expression on lifespan suggests that it may be a further target of selection, with a trade-off between increased starvation resistance but shorter lifespan associated with high *fiz* expression in females but low *fiz* expression in males. Interestingly, in females *chinmo* and *fiz* expression affected lifespan in opposing directions (Fig 5), strongly suggesting that *chinmo* expression helps modulate *fiz*’s effect on lifespan. Indeed, many of the metabolites and enriched pathways we detected are known to affect lifespan and likely contribute to *fiz*’s effect on this phenotype.

Between two genetic backgrounds in which disruption of *fiz* expression is known to drive the same phenotypes (Glaser-Schmitt and Parsch 2018), we detected only a small overlap in genes DE when *fiz* expression was disrupted (7.4–9.3% of DE genes; Table 1), with the majority of DE genes being background-specific. Interestingly, in our WGCNA, co-expression modules associated with *fiz* expression (Fig 3) were enriched for KEGG pathways and GO processes associated with phenotypes known to be associated with *fiz* expression (Fig S5, Table S6), but none of these modules included consistently DE genes. Moreover, genetic background was not significantly associated with any co-expression modules (Fig 3). Taken together, these findings suggest that outside of a few key genes, *fiz* expression contributes to a consistent final organismal phenotype via the expression of different genes in different genetic backgrounds. Thus, *fiz* may act upon several gene expression hubs and/or cascades, with the affected downstream genes changing depending on the genetic background. In keeping with this interpretation, *chinmo*, the most consistent and highly DE gene we detected, acts as an effector of JAK/STAT signaling (Flaherty et al 2010). Indeed, *chinmo* is an important developmental regulator (Zhu et al 2006, Wu et al 2012, Chafino et al 2023, reviewed in Suzuki 2026, Truman and Riddiford 2026) and is essential for maintaining the larval stage as well as correct progression through metamorphosis (Truman and Riddiford 2022). Moreover, dopamine has been shown to regulate 20-hydroxyecdysone levels (Rauschenbach et al 2007), both of which are affected by *fiz* expression (Fig 4; Glaser- Schmitt and Parsch 2018, Cavigliasso et al. 2024). Indeed, ecdysone and dopamine signaling are known to interact via a shared G-protein-coupled receptor (Srivastava et al 2005) thought to act as an integrative hub for molecular cascades during stress conditions (Petruccelli et al 2020), further underscoring that *fiz* expression likely affects various phenotypes via multiple signaling pathways.

Taken together, our results shed light on the molecular mechanisms underlying a complex case of selection on a pleiotropic gene. Our study underscores that selection does not always occur in a straightforward manner, which is why it is important to characterize individual cases of selection in natural populations. Doing so improves our understanding of how natural selection shapes complex organismal phenotypes, genetic variation, and the complex interaction between them.

## Materials and Methods

### *D. melanogaster* strains and staging

All *D. melanogaster* strains were maintained long-term under standard lab conditions (21°C, 14 hours light: 10 hours dark cycle, cornmeal-molasses media), while all larval/adult staging and lifespan assays were performed at 25°C. Flies or larvae for use in RNA-seq, metabolomics, qRT-PCR, or phenotyping assays were density controlled by placing the parents (30-50 adults in total) in cages and allowing them to lay eggs on molasses-agar plates supplemented with yeast for ∼48 hours. First instar larvae (identified based on size and mouth hook morphology) were collected and transferred either to large vials containing cornmeal-molasses medium supplemented with 0.05% bromophenol blue (Carl Roth; Karlsruhe, Germany) at a density of 200 larvae per vial for larvae or to small vials containing cornmeal-molasses medium at a density of 50 larvae per vial for adults. Late third instar wandering larvae were identified by the loss of blue color in their guts.

We obtained RNAi constructs under the control of the yeast GAL4/UAS system as well as their respective control strains containing an empty vector at the same genomic location from the Vienna Drosophila Resource Center (VDRC; www.vdrc.at; Vienna, Austria; Dietzl et al. 2007) to knockdown *fiz* (KK library; transformant ID: 107089; control ID: 60100) or *chinmo* (SH library; transformant ID: 330501; control ID: 60200) expression. Expression was knocked down by crossing control or RNAi construct males to females from either an Act5C-GAL4/2xTb driver strain, which drives ubiquitous GAL4 expression, or a Cyp6g1-GAL4 driver strain (6g1HR-GAL4-6c), which drives GAL4 expression in the larval fat body, midgut, Malpighian tubule, and gastric caecae (Chung et al. 2007) and was kindly provided by Llewllyn Green.

### RNA-seq, read mapping, and DE analysis

For RNA-seq, fat bodies of *fiz*-RNAi/Act5C-GAL4 or KK-control/Act5C-GAL4 third instar late wandering (puff stages 7-9) were dissected in cold PBS and stored in RNA/DNA shield (Zymo Research Europe; Freiburg, Germany) at -80°C until RNA extraction. To prevent potential complications from mixed sex samples, only female larvae were used for RNA-seq. RNA was extracted from 3 biological replicates of groups of 8 fat bodies using the RNeasy Plus Universal kit (Ǫiagen; Hilden, Germany) as directed by the manufacturer with the following modification: tissue was homogenized by vortexing. Poly-A selection, fragmentation, reverse transcription, library construction, and high- throughput sequencing was performed by Novogene (Martinsried, Germany) using the Illumina HiSeq 2500 platform (Illumina; San Diego, CA) with 150-bp paired reads. Additionally, 150-bp paired end data from the same sex, stage, and tissue from a population in Munich, Germany (Glaser-Schmitt and Parsch 2023) was downloaded from Gene Expression Omnibus (Accession number GSE218762, Sample accession numbers GSM6756187– GSM6756197). This dataset consisted of 5 isofemale strains: 1 strain containing a naturally-occurring *fiz* hypomorph allele and 4 control strains collected at the same location and date. The Munich dataset includes genetic variation outside of the *fiz* locus and is known to have a high level of expression variation (1,741–2,598 DE genes between any pair of strains; Glaser-Schmitt and Parsch 2023). Therefore, we analyzed it in conjunction with the RNAi lab background dataset to *i*) prevent the spurious identification of differential expression resulting from other natural genetic variation as associated with *fiz* expression, *ii*) better understand how genetic background influences the transcriptomic effects of *fiz* expression, and *iii*) identify expression differences associated with *fiz* expression that are consistent across genetic backgrounds.

We mapped reads to the *D. melanogaster* transcriptome including mRNA, rRNAs, microRNAs, and ncRNAs, with the FlyBase release version 6.65 annotation (Öztürk-Çolak et al 2024) using NextGenMap (Sedlazeck et al 2013) in paired-end mode. In downstream analyses, we included all ncRNAs but excluded rRNAs and microRNAs. Ambiguous read pairs mapping to two different genes or transcripts were discarded. The total number of read pairs per library ranged from 35.9–53.7 million, 92.4–98.8% of which could be mapped (Table S9). For differential gene expression analyses, we analyzed the sum of read counts across all of a gene’s transcripts (across all annotated exons), i.e. on the individual gene-level.

Because the two datasets were generated at different timepoints, for analyses using both datasets we first performed batch correction using ComBat-seq as implemented in the *sva* package (Zang et al 2020). To be considered as expressed in a dataset, a gene was required to have an average of at least 10 reads per sample in that dataset (170 reads total in the combined dataset and 60 reads total in the RNAi dataset). We carried out differential gene expression analysis using the negative binomial test as implemented in DESeq2 (Love et al 2014) in R (R Core Team 2022) with a 5% false discovery rate. Differential expression was detected in the RNAi dataset using a 1-factor model design with 2 levels corresponding to the *fiz*KD and controlKD strains. Differential expression was detected in the combined dataset using a 2-factor (genetic background and sample type) model design with 2 levels per factor corresponding to the *fiz*DIS or controlDIS strains for the sample type and the natural population or lab background. After initial estimation of their differential expression, *fiz* and *white*, which was used as an eye color marker in the RNAi dataset, were removed from the downstream analyses and differential expression was recalculated.

### Weighted gene co-expression network analysis

Normalized gene counts for the batch corrected combined dataset were calculated using DESeq2 (Love et al 2014). To better understand how *fiz* expression correlates with expression patterns in other genes, we performed a weighted gene co- expression network analysis (WGCNA) using the WGCNA R package (Langfelder and Horvath 2008) using the combined dataset to cluster genes into modules of genes with correlated gene expression. First, variance stabilized normalized gene counts of all analyzed genes were used to calculate a topological overlap (TOM) matrix using a power parameter of β = 17 for soft-thresholding. Gene co-expression modules were identified using hierarchical clustering and dynamic tree cutting with a minimum module size of 30 and a correlation threshold of 0.95 for module merging, which did not result in any module merging, although lower merging thresholds (as low as 0.75) also did not result in any module merging (Fig S7). For each gene expression module, the eigengenes (module summaries), which are essentially the first principal component for each module, were then correlated with *fiz* normalized gene counts and a binary approximation of the genetic background (natural population = 1 and lab background = 2) using a Pearson’s correlation and a student asymptotic *P*-value to assess significance.

### Gene set enrichment analysis

We used the clusterProfiler R package (Xu et al 2024) and a Benjamini-Hochberg (BH) multiple test correction (Benjamini and Hochberg 1995) to search for an enrichment of gene ontology (GO) biological process terms and KEGG pathways among DE genes and WGNCA expression modules.

### qRT-PCR

Fat bodies were dissected from female and male *fiz*-RNAi/Cyp6g1-GAL4 (*fiz*KD) and KK-control/Cyp6g1-GAL4 (controlKD) larvae at the late wandering third-instar stage (puff stages 7–9). Six biological replicates were prepared for each genotype and sex, each consisting of three fat bodies. Total RNA was extracted using the MasterPure RNA Purification Kit (Epicentre Biotechnologies; Madison, WI, USA) according to the manufacturer’s instructions. For each sample, 650 ng of total RNA was reverse- transcribed using LunaScript RT SuperMix (New England Biolabs GmbH; Frankfurt am Main, Germany).

Ǫuantitative PCR was performed using SsoAdvanced Universal SYBR Green Supermix (Bio-Rad Laboratories GmbH; Feldkirchen, Germany) on a ǪuantStudio 5 Real-Time PCR System (Applied Biosystems, Thermo Fisher Scientific; Waltham, MA, USA), with three technical replicates per biological replicate. Published primer sequences were used for *fiz*, *chinmo*, and *RpL32* (Pandey et al 2021; Cavigliasso et al 2024). Expression of *chinmo* was normalized to *RpL32*, and relative expression was calculated using the comparative Ct (ΔΔCt) method, with sex-matched controlKD samples as the control. Fold changes (2^^−ΔΔCt^) were plotted as means ± SEM across six biological replicates per group. Within each sex, log_2_-transformed fold changes were compared between *fiz*KD and controlKD using a two-sided Welch’s *t*-test.

### Untargeted Metabolomics

Whole female larvae of *fiz*-RNAi/Cyp6g1-GAL4 and KK-control/Cyp6g1-GAL4 were collected at the late wandering third-instar stage (puff stages 7–9). Three biological replicates were prepared for each genotype, each consisting of approximately 25 pooled larvae. Samples were immediately flash-frozen in liquid nitrogen and stored at −70°C until shipment to Novogene (Martinsried, Germany) for untargeted metabolomic analysis. Profiling was performed by UHPLC–MS/MS using an Orbitrap Exploris 120 mass spectrometer in positive and negative ion modes. Data processing, quality control, normalization, and metabolite annotation were performed by Novogene.

Metabolites were annotated using NovoMetDB, Novogene’s in-house MS/MS spectral library. Identified metabolites were categorized into 3 confidence levels, depending on the evidence for their expression. Level 1, the highest level, required matches in precursor mass (MS1), fragmentation spectrum (MS/MS), and retention time. Level 2 required matches in MS1 and MS/MS, and level 3 annotations were based on either an MS1 match alone or matches in MS1 and MS/MS to computationally predicted database entries.

Differentially abundant metabolites were identified separately for each ion mode using three criteria: a variable importance in projection (VIP) score > 1 from partial least squares discriminant analysis in metaX, a *fiz*KD-to-controlKD fold change > 1.2 or < 0.833, and a *t*-test *P* value < 0.05. KEGG pathway enrichment of differentially abundant metabolites was assessed separately for each ion mode using hypergeometric tests and a BH multiple test correction (Benjamini and Hochberg 1995). For each pathway, the proportion of differentially abundant metabolites assigned to that pathway was compared with the corresponding proportion among all detected metabolites with KEGG pathway annotations in the same ion mode. Pathways with *P* ≤ 0.05 were considered significantly enriched.

### Lifespan assay

Lifespan assays were performed using *fiz*-RNAi/Cyp6g1-GAL4 (*fiz*KD) and *chinmo*-RNAi/Cyp6g1-GAL4 (*chinmo*KD) flies, with KK-control/Cyp6g1-GAL4 (controlKD) and SH-control/Cyp6g1-GAL4 (controlSH) as the respective controls. Newly eclosed flies were collected on three consecutive days (batches A–C). Males and females were maintained together for three days, then separated into single-sex vials containing 10 females or 15 males. For each genotype, ∼10 vials of females and 8 vials of males were established, resulting in a total of 100 females and 135 males for *fiz*KD, 100 females and 136 males for controlKD, 107 females and 130 males for *chinmo*KD, and 100 females and 130 males for controlSH in our survival analysis. Flies were maintained at 25°C and transferred to fresh food every Monday, Wednesday, and Friday for the first five weeks of the assay and once weekly thereafter. Mortality was recorded every Monday, Wednesday, and Friday until all flies had died (total assay duration: 14 weeks).

Adult age was calculated separately for each collection batch. Flies recorded as escaped or accidentally killed were right-censored at the corresponding age. Survival curves were estimated using the Kaplan–Meier method. Each RNAi genotype was compared with its corresponding control using Cox proportional hazards models in R with the *survival* package (Therneau 2024). Analyses were performed separately for females and males, with genotype and collection batch included as explanatory variables. Genotype effects were assessed using a Wald test, with *P* < 0.05 considered statistically significant.

## Acknowledgements

We thank Hilde Lainer for excellent technical assistance in the lab. We thank John Parsch (JP) for providing feedback on the manuscript as well as facilities, funding, and resources. We thank Llewllyn Green for kindly providing 6g1HR-GAL4-6c flies.

